# tensorBF: an R package for Bayesian tensor factorization

**DOI:** 10.1101/097048

**Authors:** Suleiman A. Khan, Muhammad Ammad-ud-din

## Abstract

With recent advancements in measurement technologies, many multi-way and tensor datasets have started to emerge. Exploiting the natural tensor structure in the data has been shown to be advantageous for both explorative and predictive studies in several application areas of bioinformatics and computational biology. Therefore, there has subsequently arisen a need for robust and flexible tools for effectively analyzing tensor data sets. We present the R package tensorBF, which is the first R package providing Bayesian factorization of a tensor. Our package implements a generative model that automatically identifies the number of factors needed to explain the tensor, overcoming a key limitation of traditional tensor factorizations. We also recommend best practices when using tensor factorizations for both, explorative and predictive analysis with an example application on drug response dataset. The package also implements tools related to the normalization of data, informative noise priors and visualization. Availability: The package is available at https://cran.r-project.org/package=tensorBF.

## 1 Introduction

Tensor factorizations are rapidly gaining popularity in data analysis. An increasing number of tensor applications have recently emerged in bioinformatics to study diverse biological data sets. Recently, [1] applied tensor factorization to uncover novel gene networks linked to genetic variation in an *individuals x tissues x genes* tensor. Tensor factorisation has also been shown useful in predicting survival outcomes in cancers using copy number data of *individuals x genes x condition* tensor [2].

Tensor factorizations also present a novel formulation of precision medicine. Here, a natural tensor arises when the measurements of a set of biomarkers can be obtained after multiple interventions in a group of patients, yielding a *individuals x drugs x biomarker* tensor. Considering cancer cell lines as a proxy for individual patients, we formulate an application setting of tensor factorization in Fig 1. The tensor data set consists of the molecular responses (gene expression) of multiple interventions (drugs) on multiple different cancers (cell lines), obtained from the CMap [3]. Here, the key question that tensor factorisation can answer is, which parts of the drug-responses are specific to a particular cancer and which are common across various cancers. Elucidating such effects can generate hypothesis on personalised therapies, as well as increase understanding on drug action mechanisms.

**Figure 1:**
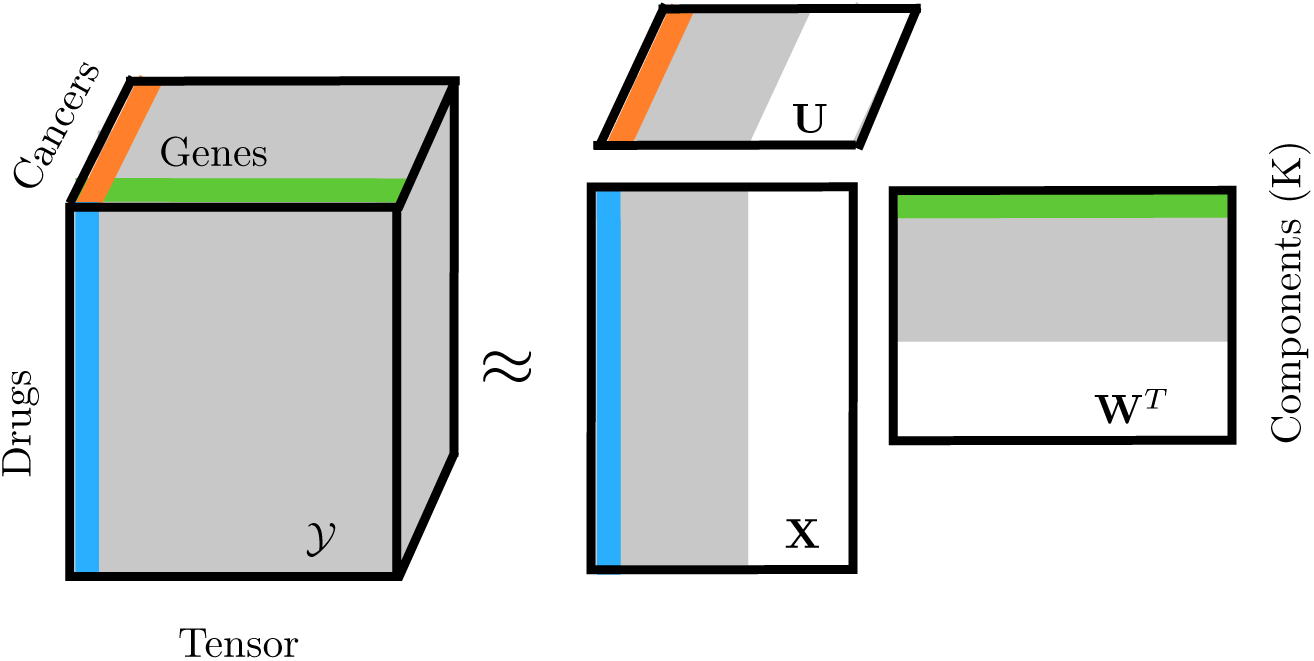
Illustration of tensor factorisation. The tensor *Y* can be factorized into a low-dimensional component space **X**,**W** and **U** which represents relationships across the drugs, genes and cancers. tensorBF automatically prunes out excessive components (shaded white in component matrices).

Fig 1 presents the well-known trilinear CP factorization of a tensor. The CP (Canonical Decomposition / Parafac) factorizes a tensor into a sum of rank-one tensors, each of which can be represented as latent variables (factors or components) in all modes [4, 5]. For the tensor ***Y*** ∊ ℝ^*N* × *D* × *L*^, CP identifies the latent variables **X** ∊ ℝ ^*N* ×*K*^, **W** ∊ ℝ ^*D* ×*K*^, and **U** ∊ℝ ^*L* ×*K*^ as

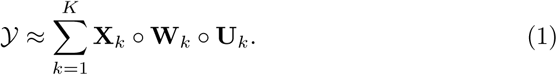

While several factorization methods exist for tensors, like the Tucker model [6], CP factorization is easier to interpret making it a promising choice for many biological applications. Recently, Bayesian tensor factorizations have been demonstrated to overcome some of the limitations including automatic determination of the number of components [1, 7], however, R package for Bayesian factorization of a tensor do not exist.

We present tensorBF, an R package to analyze natural tensor structures in the data. The package implements the Bayesian CP factorization of a tensor to infer latent factors (components) that are not obvious from the data itself. Additionally, it provides tools for analyzing the components and relationships between the variables.

## 2 Bayesian Tensor Factorization

Our package tensorBF implements the Bayesian formulation of the tensor factorization problem of Eqn (1), by assuming normal distribution with conjugate priors. A sparsity parameter is introduced that shuts down excessive components by setting them to zero (white in Fig 1), making it possible for the model to learn the true number of components automatically. Besides, the package implements feature-level sparsity for the latent variable matrices. Supplementary File 1 provides the details of the modeling assumptions and inference using Gibbs sampling.

## 3 Usage

### Model Inference and Initialization

The factorization of a 3-mode tensor *Y* can be inferred using `model <- tensorBF(Y)`, with the default options. Depending on the modeling assumptions and application setting, the function can take a variety of parameter choices as inputs. For instance, the number of components to initialize the model, how to normalize the data and an informative noise prior, that is, a user's belief on how much of the data variance should be explained with the components. A full description of the possible options is given in the functions `getDefaultOpts()` and `tensorBF()` documentation. The tensor can be normalized over different modes and ways, using `normFiberCentering()` and `normSlabScaling()`. If the features in a particular mode are deemed equally important, they should be scaled. However, if the variance is a proxy for the feature’s importance, scaling should not be done. The package manual contains simplified examples and `demo()`, demonstrating the usage of the functions on simulated data. The methods computational complexity is linear in the data dimensions and cubic only in *K*. The package took ˜1 hour for a single chain on the CMap data.

### Missing Values Prediction

The package can handle missing values by simply including them as `NA`s. The model parameters are sampled based on the observed data only, and `predictTensorBF()` predicts the missing values.

### Component Selection

The tensorBF package infers the number of components automatically. In practice, this is achieved by initializing the model with a high number of components *K* (default choice: 20% of the sum of lower two modes) and the method prunes any excessive components. The `noiseProp` in `tensorBF()` defines the proportion of variance that is expected to be explained with the components. In case, the data is expected to be heavily noisy, as with many real datasets, experimenting different choices of `noiseProp` will aid in component selection.

We explain component selection practice with a real tensor dataset of Fig 1. Fig 2 plots the methods behaviour as a function of an increasing number of initial *K*. The key observation here is that the performance improves until *K* ≤ 30, after which it stabilizes to the best result. Around the same mark, the model starts to prune all the excessive components indicating that it has already explained the data sufficiently. Therefore, in practice, we suggest to initialize *K* to a higher enough value and let the model choose the component number automatically. An appropriate *K* can be identified as one that prunes at least several excessive components.

**Figure 2:**
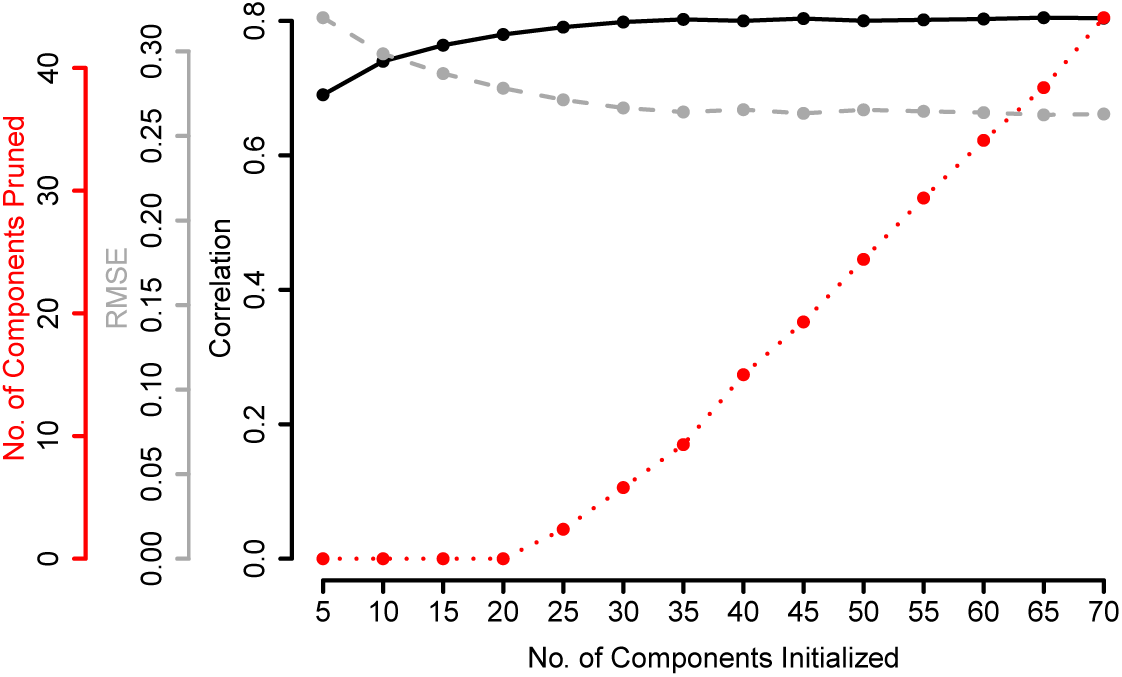
Illustrating model complexity selection with tensorBF on CMAP data set. The plot shows on the y-axis, the Pearson Correlation, Root Mean Squarred Error (RMSE), and the no. of components pruned (in red) on the missing values prediction task, as function of the number of components K used to initialize the model (the x-axis).

### Analysis and Visualizations

The factorization explains relationships between the variables through *K* components. The components can be visualized using `plotTensorBF()`. An example of such visualization is shown in Fig 3. The values of the latent variable **X** indicate that the response is primarily driven by the top 3 drugs in several HSP genes **W**. High latent scores in **U** show that this response is common across all three cancers, and can, therefore, be interpreted as a Heat Shock Protein response of HSP90 inhibitors in all three cancers.

**Figure 3:**
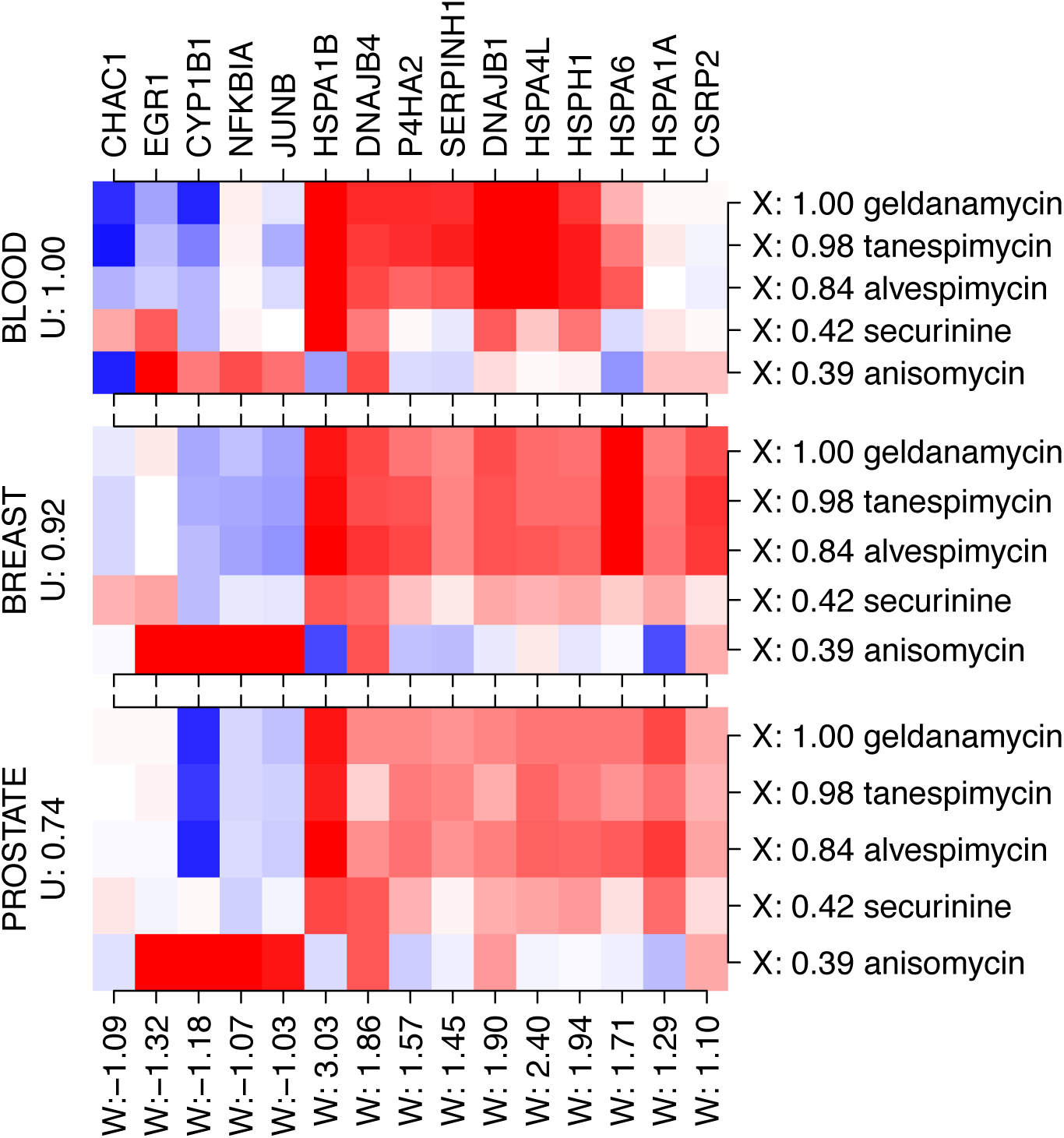
A component showing the relationship between the latent variables **X**,**W** and **U** plotted using the function *plotTensorBF().*

## 4 Discussion

The tensorBF package factorizes a tensor into low-dimensional latent factors, inferring meaningful relationships. The package provides essential tools ranging from normalization to automatic component selection, and from setting informative noise prior to interpreting the factorization. The package is a new contribution in the data analysis domain focusing on tensors with a fully Bayesian treatment of the latent factors.

